# MSclassifR: an R Package for Supervised Classification of Mass Spectra with Machine Learning Methods

**DOI:** 10.1101/2022.03.14.484252

**Authors:** Alexandre Godmer, Yahia Benzerara, Emmanuelle Varon, Nicolas Veziris, Karen Druart, Renaud Mozet, Mariette Matondo, Alexandra Aubry, Quentin Giai Gianetto

## Abstract

MSclassifR is an R package that has been specifically designed to improve the classification of mass spectra obtained from MALDI-TOF mass spectrometry. It offers a comprehensive range of functions that are focused on processing mass spectra, identifying discriminant *m/z* values, and making accurate predictions. The package introduces innovative algorithms for selecting discriminating *m/z* values and making predictions. To assess the effectiveness of these methods, extensive tests were conducted using challenging real datasets, including bacterial subspecies of the *Mycobacterium abscessus* complex, virulent and avirulent phenotypes of *Escherichia coli*, different species of Streptococci and nasal swabs from individuals infected and uninfected with SARS-CoV-2. Additionally, multiple datasets of varying sizes were created from these real datasets to evaluate the robustness of the algorithms. The results demonstrated that the Machine Learning-based pipelines in MSclassifR achieved high levels of accuracy and Kappa values. On an in-house dataset, some pipelines even achieved more than 95% mean accuracy, whereas commercial system only achieved 62% mean accuracy. Certain methods showed greater resilience to changes in dataset sizes when constructing Machine Learning-based pipelines. These simulations also helped determine the minimum sizes of training sets required to obtain reliable results. The package is freely available online, and its open-source nature encourages collaborative development, customization, and fosters innovation within the community focused on improving diagnosis based on MALDI-TOF spectra.

**Key points:**

- MSclassifR is a comprehensive R package enabling the construction of data analysis pipelines for the precise classification of mass spectra.
- Our R package contains an innovative method for variable selection from random forests, which delivered excellent results on real data.
- In-depth analysis of various machine learning-based pipelines using our package allowed us to make conclusions about the optimal m/z selection and prediction methods depending on the size of the training dataset.
- Using a publicly available dataset of mass spectra obtained from various MALDI-TOF instruments across different countries, MSclassifR is able to build robust pipelines capable of adapting to different instruments in an automatic way.
- When tested on an in-house dataset, MSclassifR pipelines consistently outperformed a commercial software in terms of prediction accuracy.

## Introduction

In clinical microbiology, the accurate identification of pathogens is crucial for proper patient management. Matrix Assisted Laser Desorption Ionization - Time of Flight (MALDI-TOF) mass spectrometry has become widely adopted for pathogen identification in biological cultures due to its speed, affordability, and user-friendly nature [1]. This technique involves comparing the mass spectrum obtained from ionizing intact proteins of a viral or bacterial culture with reference mass spectrum profiles. Commercial software provided by spectrometer manufacturers uses algorithms to match the measured mass spectrum with the spectra of known microorganisms. While this technique is effective in distinguishing many species, it often falls short in differentiating spectrally close organisms, such as subspecies within the *Mycobacterium abscessus* complex [2] [3] or species from the *Enterobacter cloacae* complex [4] [5].

To overcome these limitations, several studies have explored advanced machine learning (ML) techniques to fully exploit the information contained in mass spectra and accurately classify them into distinct categories, such as detecting resistant or virulent microorganisms in closely related mass spectra [6] [7] [8]. However, many of these studies focus on a single ML method [9] [10] [11] [12] and others lack the provision of corresponding computer code, making reproducibility of results challenging [8].

While open-source R packages like MALDIquant and MALDIrppa already offer well-documented methods for pre-processing MALDI-TOF mass spectra [13] [14], additional steps are necessary to classify the pre-processed mass spectra, including the selection of discriminative peaks and an optimal mass spectra classification model. These steps require advanced knowledge and specialized R packages for variable selection and ML methods, such as the caret and mixOmics R packages, which further demand expertise in parameter optimization [15] [16]. On the other side, non-coders have access to all-in-one software solutions, but they are often expensive, do not provide the computer code used, and provide limited user control over parameters (e.g. [17] [18]).

In response to these challenges, we have developed MSclassifR, a comprehensive package that enables the construction of data analysis pipelines for the precise classification of mass spectra. Within this article, we introduce the functionalities offered by MSclassifR and assess their effectiveness. We conducted tests using challenging real datasets and created additional datasets with varying sizes from them to evaluate the performance of data analysis pipelines built with MSclassifR.

Notably, our findings highlight the pivotal role of mass-over-charge (*m/z*) value selection in achieving accurate classifications. We have devised an innovative statistical method for variable selection, taking advantage of variable importances derived from Random Forests (RF) and employing a mixture model that distinguishes between discriminant and irrelevant features. This unique approach has led to superior performance in terms of accuracies and Cohen’s Kappa coefficient values compared to alternative feature selection methods used before a prediction model.

## Methods

The MSclassifR package offers R functions that concerns three main topics [*Figure 1A*]: (i) pre-processing of mass spectra, (ii) selection of discriminative *m/z* values, (iii) estimation of a classification model to predict the category of a mass spectrum from shortlisted *m/z* values.

**Figure 1.**
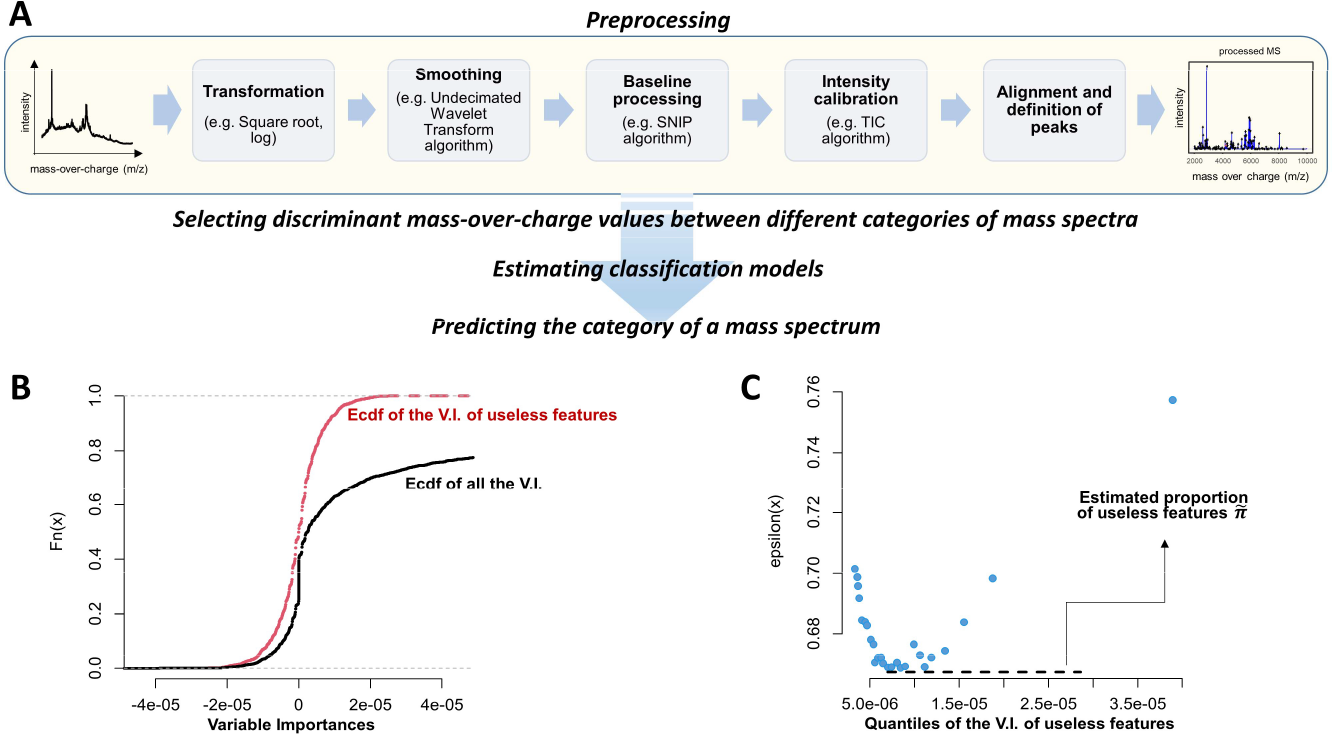
MSclassifR workflow and plots presenting the method developed for thresholding the variable importances of Random Forests. **A**. Overview of the proposed pipeline in MSclassifR, encompassing mass spectrum preprocessing with various methods, selection of discriminative *m/z*, and the estimation of a classification model to predict mass spectrum categories based on the shortlisted *m/z* values. **B**. Empirical cumulative distribution functions of the variable importances (V.I.) for the useless *m/z* (red) and all the *m/z* (black) estimated by the “SelectionVar()” function with the “mda” method from a real dataset [33] **C**. Estimation of π∼ using the same V.I. as in **B**. and the quantities ϵ_q_ = *F*(*x*_q_)/*q* where *q* are quantiles of useless variables.

### Pre-processing of mass spectra methods

Many of the algorithms already available in the MALDIquant and MALDIrppa packages are implemented in MSclassifR [13] [14]. For example, the smoothing step can be performed by using the “Savitzky Golay” algorithm or the “Undecimated Wavelet Transform” (UDWT) algorithm [19] [20] ; baseline subtraction can be performed using the “statistics-sensitive non-linear iterative peak-clipping” (SNIP) algorithm [21], the TopHat algorithm [22], or the Convexfull algorithm [23]; intensities can be normalized using the “Total Ion Current” (TIC) algorithm or the “Probabilistic Quotient Normalization” (PQN) algorithm [24]; mass spectra can be aligned using the “Locally Weighted Scatterplot Smoothing” (LOWESS) method [25] or by employing linear, quadratic, or cubic functions [26]. Finally, the peak selection step can be repeated using the “Median Absolute Deviation” (MAD) method [27] or the “super smoother” method [28]. However, we have defined a pipeline by default in MSclassifR including several algorithms that are widely used for processing MALDI-TOF MS spectra [29] [*Figure 1A*]. It consists of the following steps: (i) square root intensity transformation, (ii) spectrum smoothing with UDWT algorithm, (iii) baseline processing with SNIP algorithm, (iv) intensity calibration using TIC algorithm, (v) spectrum alignment with the LOWESS method and selection of peaks with signal-to-noise ratio (S/N).

### Selecting discriminant *m/z* values

The selection of discriminant *m/z* values is analogous to the variable selection problem in statistics. It is crucial to choose the right variables to avoid incorporating irrelevant ones into prediction models, which can lead to inaccurate predictions. Additionally, variable selection helps reduce the dataset size, which is advantageous from a computational time perspective. MSclassifR provides several methods for variable selection, which can be categorized into two groups [30]:

a. the first aims to identify a minimum number of variables that yield the highest possible performance criterion (e.g., Accuracy or Cohen’s Kappa), often resulting in the elimination of correlated variables;
b. the second aims to identify all variables with potential predictive power (e.g. without deletion of correlated variables).

On one hand, the first group is commonly employed in ML literature [30] where the primary objective is to achieve the highest accuracy with the smallest set of variables. In omics sciences, this approach is used to identify a small set of potential biomarkers (e.g. genes, proteins, metabolites) associated with a specific biological condition. A popular method of this type of approach is “Recursive Feature Elimination” (RFE) which consists of recursively selecting smaller and smaller sets of variables. This type of method typically involves setting a specific number (or range for that number) of variables to select.

On the other hand, the second group enables the filtration of variables that lack predictive power (e.g. background noise) and retains only those with predictive capabilities. For example, this approach is used to extract the set of genes or proteins that exhibit differential expression between different biological conditions. Thus, this type of method adapts to the dataset to select all the predictive variables without specifying the number of variables to select.

In addition to existing methods, we have implemented an original approach to identify all variables with predictive power using RF. This method involves estimating the distribution of variable importances for non-discriminant (useless) variables from negative importances or simulated non-discriminant variables [*Figure 1B*]. Subsequently, a mixture model is assumed, where the cumulative distribution of variable importances for all variables (*F*) is a combination of the cumulative distributions for useless variables (*F*_*u*_) and discriminant variables (*F*_*d*_), with π representing the proportion of useless variables in the dataset:

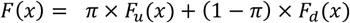

By estimating the distribution of useless variables (*F*_*u*_(*x*)), we can determine its quantile values (*x*_*q*_) and compute ϵ_*q*_ = *F*(*x*_*q*_)/*q* for a given quantile (*q*) of useless variables [*Figure 1C*]. For high quantile values of *q*, it is expected that *F*_*u*_(*x*_*q*_) tends to 1. Thus, ϵ_*q*_= π + (1 − π) × *F*_d_ (*x*_*q*_) for such quantiles when *F*_*d*_ (*x*_*q*_) is not null. It is assumed that *F*_*d*_ becomes zero when the variable importance is small. If this assumption also holds, ϵ_*q*_ tends to π. The minimum value of the ϵ_*q*_ corresponds to an estimated proportion of useless variables in the dataset, i.e. 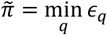 [*Figure 1C*], enabling us to estimate a number of discriminant variables (*N*_*d*_) as floor 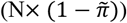, where N represents the total number of variables and “floor()” is the integer value function. Finally, the *N*_*d*_ variables with the highest variable importances are selected as the set of discriminant variables. Here, the π estimation is analogous to finding the proportion of true null hypotheses in the context of multiple hypothesis testing with the difference that we use the variable importances instead of p-values. A major difference lies in the absence of a reliable theoretical framework for estimating the distribution of variable importances for useless variables, whereas an assumption of independence between the tests and therefore a uniform distribution of p-values under the null hypothesis can be assumed in the multiple testing context [31]. It is therefore necessary to estimate the distribution of useless variables either by generating such variables or using the negative importance variables and assuming that *F*_*u*_(*x*) is a more or less symmetrical distribution with respect to zero. Both approaches are possible through our “SelectionVar()” function of our package (“*nbf”* argument). Because variable importances can be estimated with different approaches using RF, two ways of estimating them are implemented in “SelectionVar()”: “mda” for the “mean decrease in accuracy” variables importances, and “cvp” for “cross-validated permutation” variables importances [32].

### Predicting the category of a mass spectrum

Once we selected *m/z* values, a model can be estimated to predict a category from these shortlisted values. In supervised classification, a commonly used parametric model is multinomial linear logistic regression. It consists in estimating the probability that a mass spectrum belongs to a category from the following model:

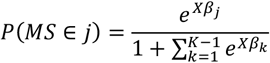

where *P*(*MS ∈ j*) is the probability that a mass spectrum belongs to the *j*^th^ category (among *K* categories), *X* are intensities of the mass spectrum measured at shortlisted *m/z* values, and *β*_j_ are coefficients related to the *j*^th^ category. Once all the coefficients are estimated, a mass spectrum is associated to a category by

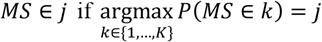

In addition to multinomial linear logistic regression, popular ML algorithms are available in MSclassifR like RF, neural networks, support vector machines with linear kernels and extreme gradient boosting. Each of these algorithms can be used to estimate a prediction model via the “LogReg()” function. Of note, the “LogReg()” function is able to deal with imbalanced datasets displaying skewed class proportions. This is a characteristic present in many real datasets when some categories are rarer than other. Different approaches are implemented to resample the data before training the algorithms via the “*Sampling*” argument: up-sampling, down-sampling and SMOTE (Synthetic Minority Oversampling Technique). Next, the “PredictLogReg()” function can be used to estimate the probabilities *P*(*MS ∈ j*) from one or several prediction models. In addition to provide the estimated *P*(*MS ∈ j*) of each model in input, “PredictLogReg()” includes two methods to aggregate the predictions of several models: the max-voting method, and the Fisher’s combined probability test to merge all the estimated *P*(*MS ∈ j*) of the models in input.

Another approach using parametric models involves conducting multiple linear regressions, one for each category, and selecting the category associated with the regression that has the lowest Akaike Information Criterion:

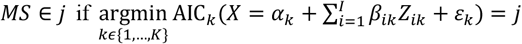

Here, *X* represents the intensities of the mass spectrum measured at *m/z* values, *Z*_*ik*_ represents the intensities of the *i*^th^ mass spectrum of the *k*^th^ category used to estimate the model at the same *m/z* values, ε_k_ denotes the Gaussian white noise corresponding to the *k*th linear regression with parameters α_*k*_ and *β*_*ik*_, and AIC_*k*_ represents the Akaike Information Criterion of the *k*th linear regression. This approach has been implemented in an original function called “PredictFastClass()” in MSclassifR. It is important to note that in order to perform the regression, the number *I* of mass spectra used within a category has to be lower than the number of *m/z* values. Additionally, for each category, a Fisher test can be conducted to determine if the *β*_*ik*_ coefficients of the linear model are zero, indicating significant correlation between the mass spectrum and at least one of the mass spectra in a category. By performing this test for all categories and selecting the minimum p-value, we can assess if the mass spectrum significantly belongs to at least one of the input categories. If the hypothesis is rejected for all performed regressions, the mass spectrum can be considered as not belonging to any of the input categories.

### Experimental datasets used to assess the performance of MSclassifR

We used MSclassifR on openly accessible datasets to assess pipelines for accurately predicting the category of a given mass spectrum. These datasets are of variable sizes:

- Dataset 1: 362 nasal swab mass spectra obtained from 312 patients categorized as either SARS-CoV-2 infected (211 mass spectra) or SARS-CoV-2 uninfected (151 mass spectra) from the Nachtigall *et al*. study [33]. These data are available at https://www.ebi.ac.uk/pride/archive/projects/PXD021388.
- Dataset 2: 882 mass spectra from 294 strains of *Escherichia coli*, classified into two categories: Shiga-toxin producer (190 strains, 570 spectra) or non-producer (104 strains, 312 spectra) from the of Christner *et al*. study [6]. These data are available at https://zenodo.org/record/4996418.
- Dataset 3 (in-house): 1001 mass spectra from 41 clinical strains representing the three subspecies of the *Mycobacterium abscessus* complex: *M*. subsp. *abscessus* (15 strains, 633 spectra), *M*. subsp. *bolletii* (7 strains, 164 spectra), and *M*. subsp. *massiliense* (9 strains, 201 spectra). These data are available at https://zenodo.org/record/5793313.
- Dataset 4 (in-house): 1890 mass spectra from 79 clinical strains from *Streptococcus pneumoniae* (60 strains, 1434 spectra) and from the viridans streptococci group including *S. mitis* (11 strains, 264 spectra) and *S. pseudopneumoniae* (8 strains, 192 spectra). These data are available at https://zenodo.org/record/8419282.

### General design of numerical experiments

In our experiments, we employed the previously discussed datasets and used the mass spectra from each dataset where we separated each into a training set of mass spectra, consisting of 70% of randomly chosen mass spectra, and we used the rest of the mass spectra (30%) to test the prediction models and evaluate the accuracy and the Kappa values. This was repeatedly performed several times to assess the variations of accuracy and Kappa values. For the training datasets, pre-processing of mass spectra was performed similarly on all datasets with the “SignalProcessing()” function of our R package by the following steps: (1) transforming intensities with sqrt method; (2) smoothing with UDWT algorithm; (3) remove baseline with SNIP method; (4) calibrate intensity with TIC method; (5) aligning spectra with LOWESS method and with a tolerance parameter of 5e-04 ppm. Next, intensity peaks of mass spectra were detected using the “PeakDectection()” function of our R package with default parameters (tolerance parameter of 2e-03 ppm, signal-to-noise ratio (SNR) of 3, and halfWindowSize parameter of 11). Mass spectra were normalized according to the maximum intensity value for each mass spectrum. For the testing datasets, the same pre-processing of mass spectra was used except that they were aligned on a reference mass spectrum created from the ones of the training dataset using a tolerance parameter of 5e-04 ppm, a minFrequency parameter of 70% and the “strict” method for the “*binPeaks_method*” argument of the “SignalProcessing()” function. Indeed, the alignment to a reference mass spectrum is important to ensure that the mass spectra in the testing dataset closely match those used during the training of the prediction model.

We conducted extensive comparisons by employing the previously discussed datasets 2 to 4 and generating subsets of randomly chosen mass spectra of various sizes among them. Each subset has been created by respecting the original proportions of each category in the datasets. To avoid overfitting, we ensured that when multiple replicates of a particular sample were present, they were consistently grouped together within the same subset. Here, our main focus revolved around the training dataset sizes and the advantages of using contemporary ML methods (RF, neural networks, support vector machines with linear kernels and extreme gradient boosting) compared to conventional parametric approaches such as decision rules based on multinomial linear logistic regression or multiple linear regressions, as presented earlier. Additionally, it is important to evaluate the number of mass spectra needed to construct an accurate classifier.

## Results and discussion

### Extensive comparison from randomly generated subsets of mass spectra

For the datasets (2 to 4) previously described, we randomly drew sets of mass spectra of different sizes. These generated datasets range from 50 mass spectra to 1890 mass spectra. For each of them, a training set consisting of 70% of the mass spectra from the generated dataset and a test set (composed of the remaining 30% of mass spectra) were constituted several times by random draws. For each generated dataset, we evaluated 6 different methods available in MSclassifR to select discriminative peaks, which we combined with 6 different methods to predict the different categories of mass spectra on the test sets. All the parameters used for the various functions are in table S4. The 36 pipelines were assessed on the test datasets, using both Accuracy and Cohen’s Kappa criteria, leading to 8604 assessments (Table S1).

Using this methodology, we have made a number of observations on (i) the impact of the *m/z* selection method on the performances of classification models (ii) the impact of dataset size on the selection of *m/z* values and (iii) the global performances of the used pipelines (*m/z* selection + prediction algorithms).

The first observation (i) suggests the importance of performing a variable selection step before estimating a classification model. Regardless of the prediction algorithm used and the dataset size, the “cvp” and “mda” *m/z* selection methods significantly improve both Accuracy and Cohen’s Kappa when compared to not selecting *m/z* (e.g. estimating the classification model on all the *m/z*) or using the “SelectionVarStat”, “RFERF”, or “sPLSDA” methods [*Figure 2A*]. This observation must nevertheless be put into perspective on data sets composed of many mass spectra (more than 1000). In such cases, “sPLSDA” and “RFERF” demonstrate similar performance levels to “cvp” and “mda”, while the “SelectionVarStat” method’s performance declines with increasing dataset size, leading to performances similar to not selecting *m/z* values [*Figure 2A*].

**Figure 2.**
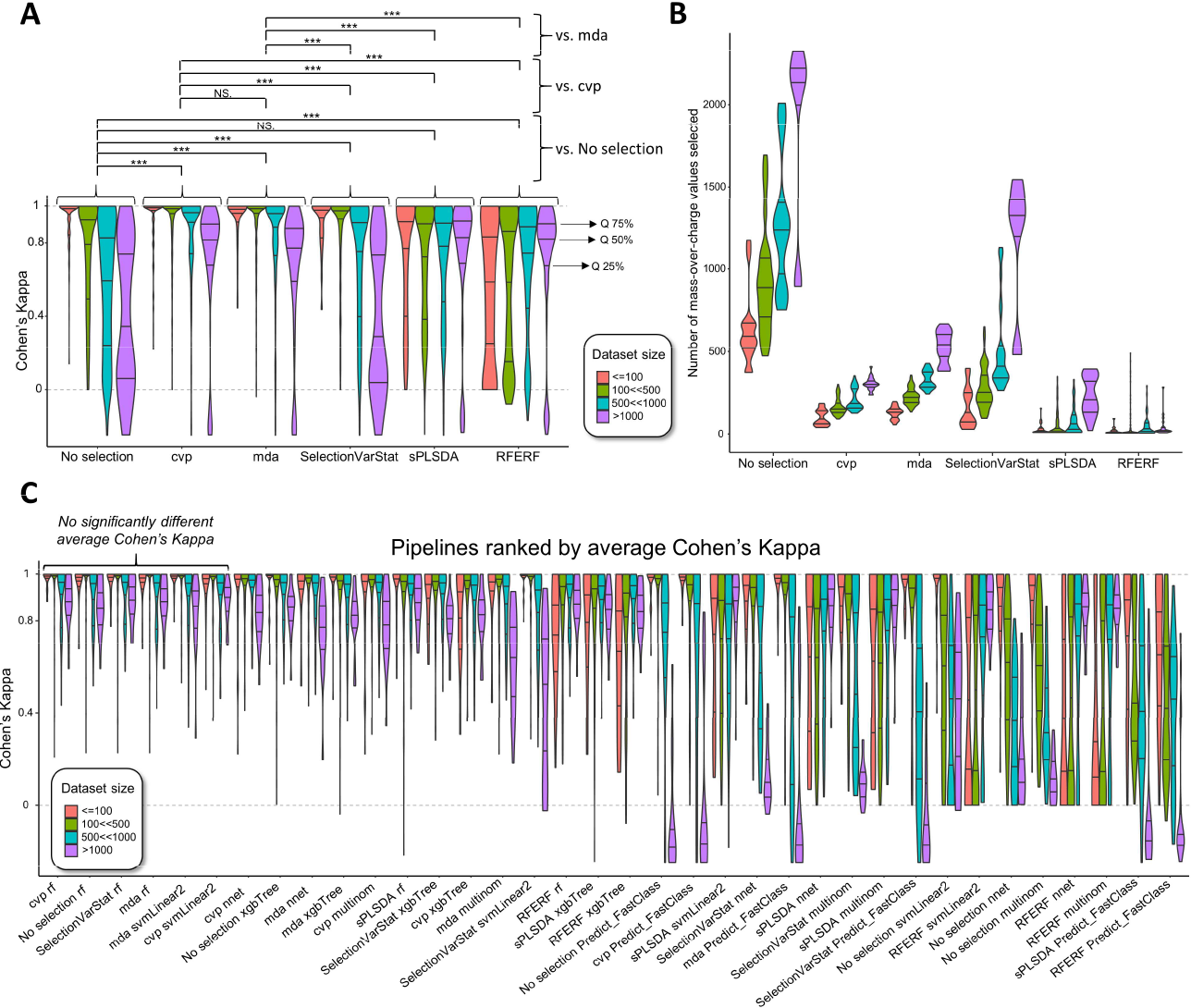
Performances of pipelines implemented in MSclassifR on 239 datasets of various sizes. **A**. Violin plots of Cohen’s Kappa depending on the variable section method used. The dataset sizes are in different colors. **B**. Violin plots of the number of selected *m/z* depending on the variable section method used. The dataset sizes are in different colors. **C**. Violin plots of the Cohen’s Kappa depending on the pipeline (variable selection + prediction algorithm) used. Pipelines are ranked from the highest average Cohen’s Kappa (left) to the lowest (right) estimated on all the generated datasets. The dataset sizes are in different colors. **Notes:** cvp: for cross-validated permutation variables importances; mda: mean decrease in accuracy; multinom: logistic regression; nnet: single-hidden-layer neural network; No selection: no *m/z* selection method; Predict_FastClass: multiple regression estimator; RFE-RF: Recursive Feature Elimination with Random Forests; SelectionVarStat: ANOVA test; RF: random forests; sPLSDA: Sparse Partial Least Squares Discriminant Analysis;; svmLinear2: linear Support Vector; xgbTree: eXtreme Gradient Boosting. * *p-value* < 5.10^−2^, ** *p-value* < 10^−2^, *** *p-value* < 10^−3^ (estimated using univariate Wilcoxon rank sum test).

The second observation (ii) pertains to the number of selected *m/z* values. When there are more mass spectra in a training dataset, the selection methods tend to select a greater number of potentially discriminant *m/z* [*Figure 2B*]. In general, “SelectionVarStat” tends to select the most *m/z* values among the different methods, followed by “mda” and “cvp”, while “sPLSDA” and “RFERF” tend to select very few *m/z* compared to the other methods [*Figure 2B*]. Selecting more *m/z* can potentially lead to reduced performance due to errors in mass spectra classification as previously observed with “SelectionVarStat”. That is why “sPLSDA” and “RFERF” are relatively robust to the dataset size increasing [*Figure 2A*].

Concerning the global performance of the MSclassifR pipelines (observation (iii)), RF globally outperform other prediction models [*Figure 2C*]. When assessing the effectiveness of variable selection methods in combination with RF, “sPLSDA” and “RFERF” exhibit significantly lower accuracy and Cohen’s Kappa values compared to alternative methods [*Figure 2C*]. Importantly, there is no significant difference in performance when combining RF with the other methods “cvp,” “mda”, “SelectionVarStat”, and “No selection”. These similar performances can be attributed to the inherent functioning of the RF algorithm: it employs a random variable selection process during the construction of each decision tree that can make the selection made upstream obsolete. Interestingly, after performing *m/z* selection with “mda” or “cvp”, support vector machines with linear kernels “svmLinear2” perform similarly to RF and could also be an interesting strategy to apply [*Figure 2C*]. Thus, using “mda” or “cvp” *m/z* selection methods combined with either “rf” or “svmLinear2” often deliver some of the best performances across different dataset sizes, with no significant differences between them. We have also observed that certain strategies perform well on small size datasets but exhibit poor performance on larger ones. Specifically, “Predict_FastClass” associated with “mda”, “cvp”, “SelectionVarStat” or “No selection”, as well as “svmLinear2” combined with “SelectionVarStat” or “No selection”, fall into this category [*Figure 2C*]. While these strategies exhibit competitiveness when confronted with constrained numbers of mass spectra in a training dataset, their performance should be approached cautiously in broader contexts.

### Application on another independent dataset

We aimed to determine if the promising pipelines, identified through extensive comparisons, could perform well on a separate dataset (Dataset 1) with its own unique challenges [33]. This particular dataset presents the added complexity of spectra obtained from various MALDI-TOF instruments across different countries. It is worth noting that the original authors applied a calibration function to mitigate inter-laboratory differences in mass spectra, a step we did not replicate in our computational experiments.

In our experiments, we applied the same preprocessing steps to the mass spectra as before. We then rigorously evaluated the models by randomly dividing the spectra into 70% for training and 30% for testing. This process was repeated 15 times, resulting in an average Accuracy of 0.875 and a maximum of 0.925 for the “mda rf” pipeline, which had previously shown good performance (Table S2). Notably, these precision levels closely match those reported in the article related to the dataset, even though we did not perform any calibration. Thus, it shows the ability of MSclassifR to build robust pipelines capable of adapting to different instruments in an automatic way.

To get the most out of MSclassifR, we assessed its capacity to combine prediction probabilities through ensemble-learning methods like maximum voting and Fisher’s method. Simultaneously, we examined the potential enhancements that MSclassifR’s resampling techniques could bring to our results. For this purpose, we derived five models from the “mda rf” pipeline and merged them after the application of diverse dataset resampling approaches. Our findings reveal that employing up-sampling and merging with either the Fisher’s method or the maximum vote technique leads to a significant improvement in overall performance on average (average Accuracy of 0.895 and 0.894 respectively, with maximums of 0.926 and 0.935) [*Figure 3A*]. In contrast, the down-sampling method yielded inferior results when compared to the absence of resampling [*Figure 3B*]. In addition, these ensemble-learning approaches make it possible to reduce the variability of Accuracy, and therefore to obtain more robust results [*Figure 3A-C*].

**Figure 3.**
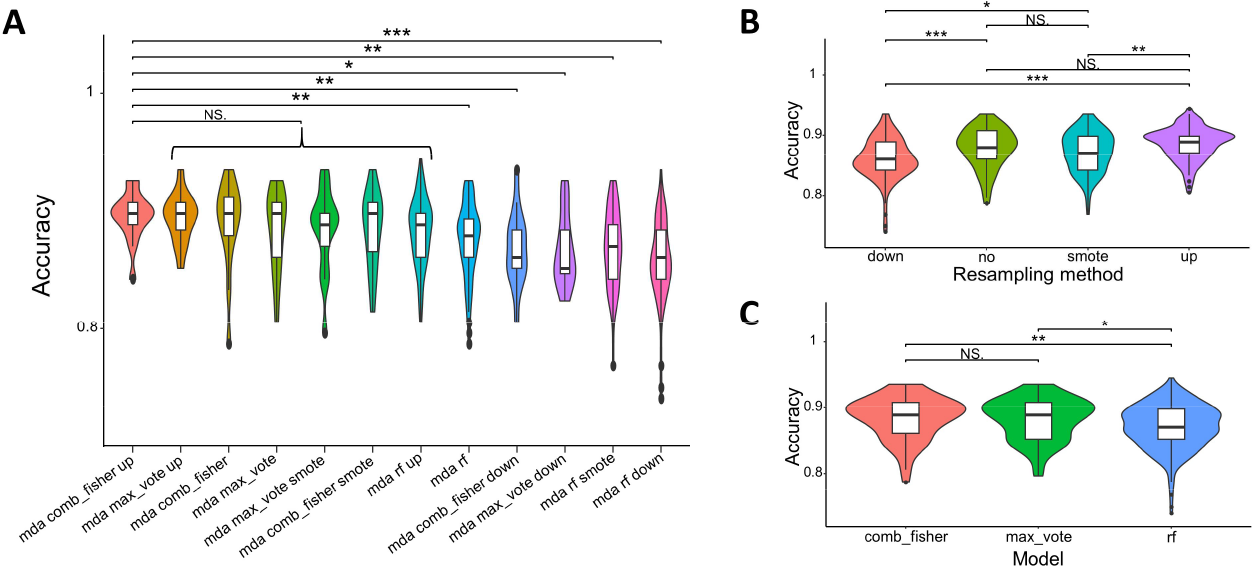
Performances of pipelines implemented in MSclassifR on dataset 1 using 70% of the mass spectra to train the pipelines and 30% to test (15 repeats for each). **A**. Violin plots of the Accuracy depending on the pipelines used on Dataset 1. **B**. Violin plots of the Accuracy depending on the resampling method used. **C**. Violin plots of the Accuracy depending on the merging technique used. * *p-value* < 5.10^−2^, ** *p-value* < 10^−2^, *** *p-value* < 10^−3^ (estimated using univariate Wilcoxon rank sum test). **Notes:** comb_fisher: combine predictions with Fisher’s method; down: down resampling method; max_vote: combine predictions with hard voting system; mda: mean decrease in accuracy; RF: random forests; up: up-resampling method; smote: SMOTE resampling method.

### Comparison with a commercial MALDI-TOF MS system

Finally, we also conducted a comparison between all the pipelines within MSclassifR using dataset 3 [34] and a commercial MALDI-TOF MS system (Microflex^®^ LT instrument (Bruker^®^ Daltonics) IVD MALDI Biotyper^®^). This dataset comprises mass spectra from three genetically narrowly related subspecies of the *Mycobacterium abscessus* complex. The data set is particularly challenging because these subspecies are spectrally very close and difficult to distinguish. The subspecies were characterized using commercial molecular methods [34]. The commercial system provided subspecies information for only 60% of the mass spectra, with a correct identification rate of 62% (Table S3). In comparison, using only 70% of all the 1001 mass spectra to train the pipelines and 30% to test them, this is randomly repeated 5 times, the pipelines of MSclassifR provided a subspecies for all the mass spectra of the test set. The best average accuracy was achieved by the “sPLSDA rf” pipeline, with an average accuracy of 96.4% and a standard deviation of 0.029 from 157.4 selected m/z values in average (Table S1). This highlights the ability of MSclassifR to achieve robust classification accuracy, overcoming some limitations of commercial systems.

## Conclusion

MSclassifR is a freely available R package, accessible via the CRAN repository network. It is designed for the automatic classification of mass spectra into predefined categories. In this paper, we have rigorously tested the capabilities of our package on datasets acquired from various sources and instruments, including both publicly available data and MALDI-TOF mass spectra obtained in-house. The results from our investigation reveal that MSclassifR can yield prediction models with excellent accuracy and Kappa values. Furthermore, when we compared MSclassifR predictions to those generated by commercial software on in-house dataset, we observed significant performance advantages for MSclassifR prediction models. Our article also delves into an extensive exploration of diverse pipelines for constructing prediction models with our package. From this exploration, we have drawn some important conclusions:

a. A selection of discriminant *m/z* values prior to the estimation of the prediction model makes it possible to improve the accuracy of predictions.
b. The original selection method that we developed and based on thresholding the variable importances of RF leads to better accuracy in predictions, whatever the size of the training dataset.
c. Overall, the prediction method providing the best average accuracies on the used datasets is the RF algorithm.
d. Some *m/z* selection methods perform poorly on small size datasets (less than 100 mass spectra) and provide good performances on large datasets (more than 1000 mass spectra).
e. Some prediction methods provide good performances on small size datasets (less than 100 mass spectra) but perform badly on large datasets (more than 1000 mass spectra).
f. Using ensemble learning and resampling techniques can provide better and more robust performances.

These outcomes are specific to the datasets and the function parameters we used, and their general applicability to any dataset might vary. Nonetheless, they provide valuable insights into creating reliable prediction models from new coming mass spectra datasets.

It should be emphasized that MSclassifR can be used in broader contexts than MALDI-TOF mass spectrometry, for example when differentially analyzing quantified values of analytes (e.g. genes, proteins, etc.) between biological conditions. Notably, MSclassifR can be used for the analysis of liquid chromatography coupled to tandem mass spectrometry (LC-MS/MS) data [35]. This technique can also help accurately distinguish closely related microbial variants, which could promote a deeper understanding of resistance mechanisms to antibiotics [36].

In conclusion, MSclassifR offers an attractive solution for biology laboratories seeking to identify pathogens from mass spectra. Its open-source nature fosters collaboration, customization, and innovation within the community dedicated to enhancing diagnostic capabilities using mass spectra. We have provided online vignettes that demonstrate how to use the package effectively. We are committed to regular maintenance and continuous improvement of our package.

## Data and availability

The MSclassifR R package is available online on the CRAN: https://cran.r-project.org/web/packages/MSclassifR/index.html

Links towards different vignettes illustrating how to use the functions of this package from real data sets are also available from this webpage.

All datasets used in this paper are available online on PRIDE and zenodo websites.

## Authors’ contributions

A.G. and Q.GG. conceived the project. A.G. and Q.GG. designed the algorithms and developed all the programs. M.M., A.A., N.V., E.V. and Y.B. have participated in design of the study and coordination and helped to draft the manuscript. A.G., Q.GG. and K.D performed computational work and analysed the data. All the authors reviewed the manuscript. All authors read and approved the final manuscript.

